# Inhibition of MRN activity by a telomere protein motif

**DOI:** 10.1101/2021.03.30.437761

**Authors:** Freddy Khayat, Elda Cannavo, Majedh Alshmery, William R. Foster, Charly Chahwan, Martino Maddalena, Christopher Smith, Anthony W. Oliver, Adam Watson, Antony M. Carr, Petr Cejka, Alessandro Bianchi

## Abstract

The MRN complex (MRX in *Saccharomyces cerevisiae*) initiates the repair of DNA double-stranded breaks (DSBs) and activates the Tel1/ATM kinase, which orchestrates the DNA damage response (DDR). Telomeres prevent DDR activation at chromosome ends, partly by keeping MRN-ATM in check. We show that the multiple activities of the MRX complex are disabled by telomeric protein Rif2 through the action of a short motif (MIN, <u>M</u>RN/X-<u>in</u>hibitory motif) at the N-terminal end of the protein. MIN executes telomeric suppression of Tel1, DDR and and non-homologous end joining (NHEJ) via direct biding to the N-terminal region of Rad50. A combination of biochemical and genetic data suggests that Rif2 promotes a transition within the MRX complex that is not conductive for endonuclease activity, DNA-end tethering or Tel1 kinase activation. We suggests that the MIN motif operates in the *RIF2* paralog *ORC4* (Origin Recognition Complex 4) in *K. lactis* and in telomeric protein Taz1 in *Schizoccharomyces pombe*, which is not evolutionarily related to Orc4/Rif2. These results highlight a potential Achilles’ heel in Rad50, the regulatory subunit of MRN, which we suggest has been targeted by different telomeric factors in multiple fungal lineages, raising the possibility that analogous approaches might be deployed in other Eukaryotes as well.

## Introduction

Telomeres make a crucial contribution to genome stability by ensuring both the full replication of the ends, through the action of the telomerase enzyme (reviewed in ^1^), and their protection, through the agency of the telomeric complex (shelterin in vertebrates, reviewed in ^2^). Telomerase synthesises short DNA repeats onto chromosome ends, which serve as a platform for the binding of telomere-specific and accessory factors. These factors, in turn, modulate the activity of the telomerase enzyme, and the feed-back at each telomere determines the ultimate length of the telomeric DNA repeat array, which is mostly present in double stranded form but terminates in a single-stranded 3’ overhang. Although it was early on predicted that telomeres would need to be equipped with mechanisms to disable the various arms of the DNA damage response (DDR) machinery, it was perhaps unanticipated that several of the DDR factors would be instrumental in ensuring telomere end processing and telomerase activity, as first documented in budding and fission yeast ^3–6^. While several differences in the way the telomeric complex interacts with and affects DDR factors in different organisms are well documented, several common themes have emerged (reviewed in ^7,8^). An example where elements of the DDR have been co-opted for telomere function is in the activation of telomerase. In the buddying yeast *Saccharomyces cerevisiae*, the MRX complex, which is composed of Mre11, Rad50 and Xrs2 (Nbs1 in most other organisms) and is responsible for the recruitment/activation of the DDR kinase Tel1 (the ATM ortholog in yeast), promotes telomerase action: ablation of any of the three subunits, or of Tel1, leads to dramatic telomere shortening ^3,9,10^. Similar requirements for the Tel1/ATM or the ATR (Rad3 in fission yeast) kinase for telomerase activation take place in mammalian cells and in fission yeast ^11–13^.

The MRN complex is instrumental in orchestrating both DDR signalling and DNA repair, and acts as the main sensor of double-stand breaks (DSBs) ^14^. The complex is equipped with nuclease and ATPase activity by Mre11 and Rad50 subunits, respectively, and has some structural similarity to the SMC family of proteins, with elongated coiled coil (CC) motifs from the Rad50 subunits which fold back onto themselves at a Zn-hook formed at a CXXC motif found at the centre of the coiled region in Rad50. At the base of the CC region a globular domain is formed where ATP binding cassettes from two molecules of Rad50 come together and join an Mre11 dimer, in a tetrameric assembly. Nbs1/Xrs2 contains several protein interaction modules, and is responsible for binding the Tel1/ATM kinase ^15–17^. ATP binding and hydrolysis by Rad50 lead to conformational changes within the complex, and it has been proposed that the transition from an ATP-bound ‘closed’ complex to an ‘open’ one after ATP hydrolysis regulates the multiple activities of MRN ^18–21^. The ATP-bound ‘closed’ complex is required for the tethering of DNA ends, which promotes one of the two main pathways for the repair of DSBs, non-homologous end joining (NHEJ) ^20^. MRX/MRN stimulates end joining both in budding and fission yeast ^22,23^. The ATP binding and hydrolysis by Rad50 are also required to activate Tel1/ATM ^24,25^. On the other hand, the ‘open’ state that results from ATP hydrolysis is competent for DNA binding by Mre11 and nucleolytic action, whereas Rad50 in the ATP-bound state blocks access of the nuclease active sites ^26,27^. The Sae2 factor is required for the endonuclease action of Mre11 ^28^. Endonucleolytic cleavage also appears to require ATP hydrolysis and not just release from the ‘closed’ state, so it has been proposed that an intermediate state between ‘closed’ and ‘open’, yet to be observed, might exist to promote endonucleolytic action ^14^. The endonuclease action of Mre11 is followed by an exonucleolytic one to resect DNA and help channel DSB repair away from NHEJ and toward homology-dependent repair (HDR) ^29,30^.

Telomeres must deal with the multiple threats posed by MRN in potentially initiating NHEJ, HDR, or DDR signalling. Mammalian telomeres engage the telomere binding protein TRF2 to suppress ATM signalling, presumably via formation of t-loop structures, and to directly bind Nbs1 to modulate repair outcomes at telomeres ^30,31^. In budding yeast, a different mechanism for MRX control has been described which relies on telomere protein Rif2. Rif2 is unique to budding yeast and closely related species (see below) and interacts with the C-terminal domain (RCT) of the main DNA binding protein of budding yeast telomeres, Rap1, for its association with telomeres, where it suppresses telomere elongation and NHEJ ^32,33^. The first clues that Rif2 is linked to the regulation of MRX-Tel1 came from epistasis analysis of telomere length and from a screen for suppressors of hydroxyurea sensitivity in Tel1-overexpressing cells, which produced Rif2 ^34,35^. Rif2 inhibits MRX-dependent resection in G1 ^36^ and the tethering of DNA ends after DSB formation ^23^. Interestingly, the binding of MRX to DSB is not severely diminished in the absence of Rif2, suggesting a regulatory role for Rif2 ^23,37^, an idea which is supported by the observation that Rif2 affects the ATPase activity of Rad50 ^23^.

We focus here on a conserved motif within Rif2, previously shown to be involved in the regulation of telomere length and Tel1 activation ^38,39^, which we refer to as MIN for <u>M</u>RN-<u>in</u>hibitor. We show, by a combination of genetic and biochemical approaches, that the motif is a potent inactivator of MRX-Tel1, as it is capable of disabling the action of MRN-Tel1 in telomerase activation, NEHJ suppression, and endonucleolytic digestion of DNA. Our data are consistent with a direct action of the motif on the N-terminus of Rad50, and an effect on the ATP-hydrolysis-dependent Rad50-mediated allosteric transition enacted by Rad50 on the complex. We suggest that MIN provides a powerful and comprehensive mechanism for telomeres to inactivate the multiple actions of MRN-Tel1 at chromosome ends, and that this mechanism has independently arisen in multiple lineages during fungal evolution.

## Results and Discussion

### Identification of a conserved motif in Rif2 and Orc4

*RIF2* evolved from the gene coding for the Origin Recognition Complex 4 subunit (ORC4) after a whole genome duplication (WGD) event that occurred in budding yeast and related species (see Figure 1e for an illustration of evolutionary relationships of species discussed in this study; see also Figure 5a, for an illustration of the WGD event within the Saccharomycotina) ^33,40^. Therefore, to gain insight into the parts of Rif2 of possible functional significance (Figure 1d), we decided to carry out an analysis of the Rif2 and Orc4 amino acid sequence aimed at identifying conserved and diverging regions. All yeast species bearing *RIF2* also carry a copy of *ORC4*, the latter being an essential gene. Clustal Omega alignment of Rif2 and Orc4 protein sequences from several fungal species in the Saccharomycetceae clade revealed that Rif2, compared to its Orc4 counterpart, consistently bore an extra stretch of about 30 amino acid at the N-terminus (Figure 1a). Strikingly, a survey of Orc4 sequences in other Saccharomycotina species devoid of Rif2, revealed that several Orc4 proteins share homology at their N-terminus to the N-terminal ‘tail’ region in Rif2 (Figure 1b). A strongly conserved consensus motif can be derived from this analysis and is summarised in Figure 1c. Four positions in particular appear to be invariantly conserved and correspond to position 7 (D/E), 8 (F), 11 (ILMV) and 12 (KR) of *Saccharomyces cerevisiae* Rif2, and we propose that these residues (in blue in Figure 1c) delimit a core motif of functional significance (see below). In this manuscript, we will refer to this motif as the MIN motif, for <u>M</u>RX-<u>in</u>hibitory motif, based on its proposed function.

**Figure 1.**
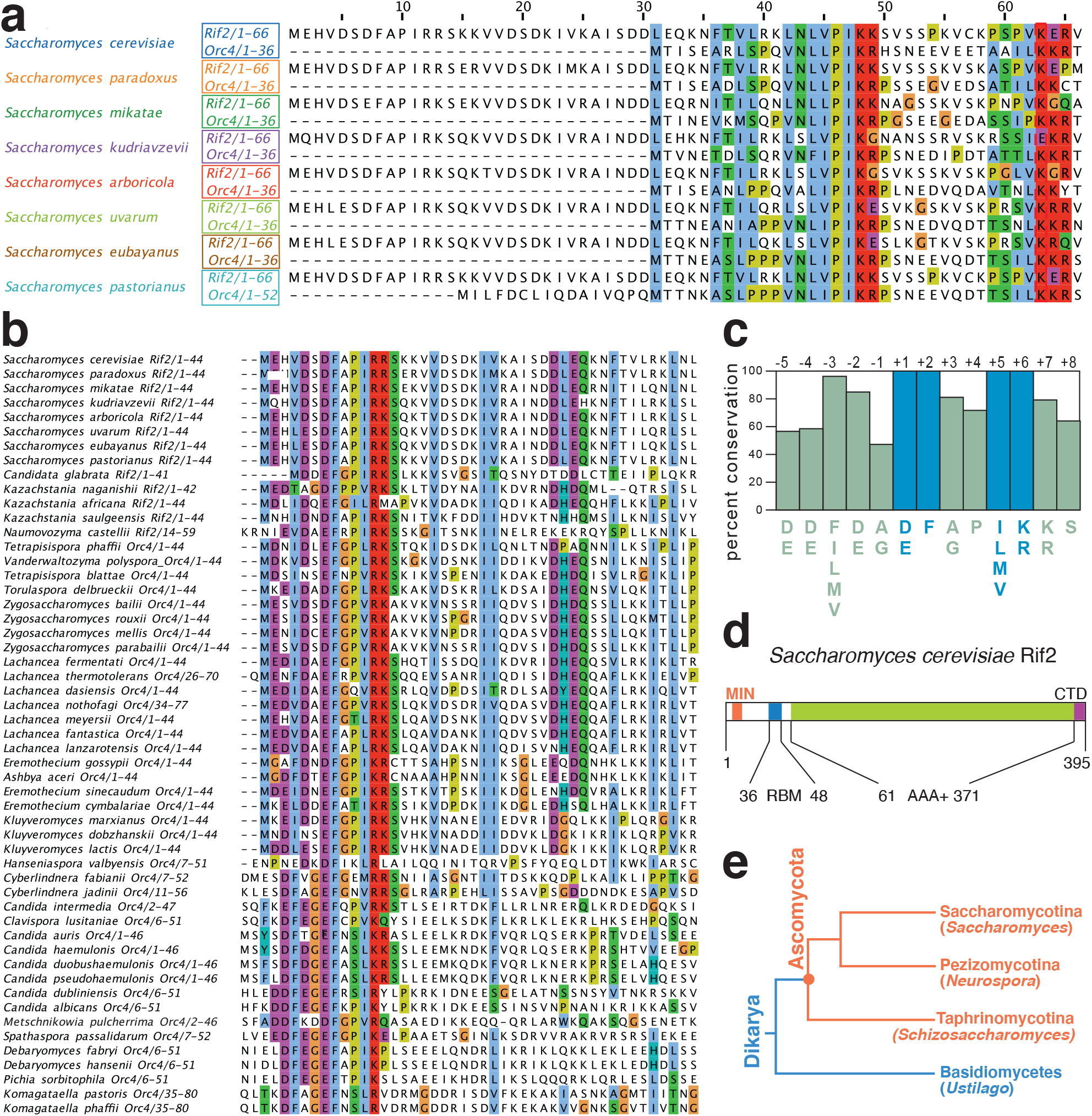
A conserved protein motif at the N-terminus of Rif2 and Orc4 in the Saccharomycotina. A) Sequence alignments of the indicated Rif2 and Orc4 proteins from Saccharomycotina species where Rif2 is present were produced in Clustal Omega and gaps were manually eliminated. Amino acid included are indicated after the protein name. Alignments were done using the Clustal Omega interface on the EBI site. Alignments were edited (in this case) and imaged in Jailview using Clustalx colouring of conserved residues (Figures 1A, 1B, 6A). B) Sequence analysis as in (A) of Rif2 and Orc4 N-terminal regions from species where the conserved region identified in the N-terminal extension in (A) is present. Species are broadly arranged top to bottom according to the phylogeny described in ^73^. C) Diagram illustrating the prevalence of conserved residues in the N-terminal cluster derived from the alignment in (B), defining the MIN motif. Based on the presence of 4 invariant residues (see also Figure 7B) we adopt for the motif the numbering indicated on top of the diagram (invariant positions at +1, +2, +5, and +6, in blue). D) Diagram of the domain structure of budding yeast Rif2. Conserved domains previously identified in ScRif2 include an AAA+ fold shared with Orc4, and a Rap1 binding motif (RBM) at the N-terminus and one (CTD, C-terminal domain) at the C-terminus ^61^; the position of the MIN motif is shown in orange. E) Phylogenetic tree, showing the relationship among the three clades comprising the Ascomycota. Representative species are indicated in parenthesis. Adapted from ^74^.

### The MIN motif regulates telomere length through the MRX-Tel1 pathway

Previous scanning mutagenesis analysis of Rif2 revealed that a small N-terminal region in Rif2 is responsible for regulating telomere length in budding yeast and defined the first 60 amino acids of ScRif2 as the BAT domain, for Blocks Addition of Telomeres ^38^. The BAT domain fully contains the MIN motif and the mutagenesis investigation of BAT is broadly consistent with the analysis of sequence conservation (Figure 1b), in that some of the most strongly conserved positions within the Sc MIN (particularly F8) were found to have the strongest telomere length phenotype when changed to alanine ^38^. Intriguingly, BAT lays within a small N-terminal fragment previously proposed to interact with the C-terminus of Xrs2 and to compete for its binding to Tel1 ^37^. This raised the possibility that MIN might function by promoting binding of Rif2 to Xrs2 and thus disabling its critical function in the Tel1 activation of telomerase. Although earlier experiments concluded that BAT does not act by affecting the function of the Xrs2 C-terminus in telomere length regulation ^38^, this analysis was carried out with an Xrs2 allele only partly defective for Tel1 binding, and showing only a mild telomere shortening phenotype ^41^. Xrs2 carries two putative Tel1-interacting motifs at its C-terminus and impairment of both motifs is required to confer significant telomere shortening ^16,38,42^. In order to determine whether the Rif2 MIN motif acts through the MRX-Tel1 pathway of telomerase regulation, we carried out epistasis analysis of telomere length using *xrs2* and *tel1* null alleles, as well a C-terminal truncation of Xrs2 which ablates both Tel1-interacting motifs in the protein and fully disables Tel1 recruitment to telomeres ^42,43^. This allele, *xrs2-664*, confers telomeres as short as the *xrs2* or *tel1* null alleles and is therefore an ideal choice for epistasis analysis (Figure 2a). As expected, a *rif2* allele carrying two point-mutations within the MIN motif (F8E and P10N mutations, *rif2-min* allele) generated telomeres at least as long as the *rif2* null. An examination of telomere length in the double mutants carrying the null allele pairings *rif2-Δ xrs2-Δ* and *rif2-Δ tel1-Δ* showed, as anticipated based on the previously recognised role of Rif2 in modulating MRX-Tel1 function ^35,44^, a dramatic shortening of telomere length near the levels seen in the *xrs2-Δ* and *tel1-Δ* single mutant strains. Importantly, we observed similarly shortened telomeres in double mutant configurations where the *rif2*-*Δ allele* was replaced by the *rif2-min* allele, suggesting that the Xrs2 C-terminus and the Tel1 kinase are both required for the telomere elongation effect conferred by loss of the MIN motif in Rif2. Thus, as the MIN motif is epistatic to Xrs2/Tel1, the simplest interpretation of these results is that MIN acts by impairing the pathway of telomerase activation that relies on the MRX-dependent recruitment of the Tel1 kinase.

**Figure 2.**
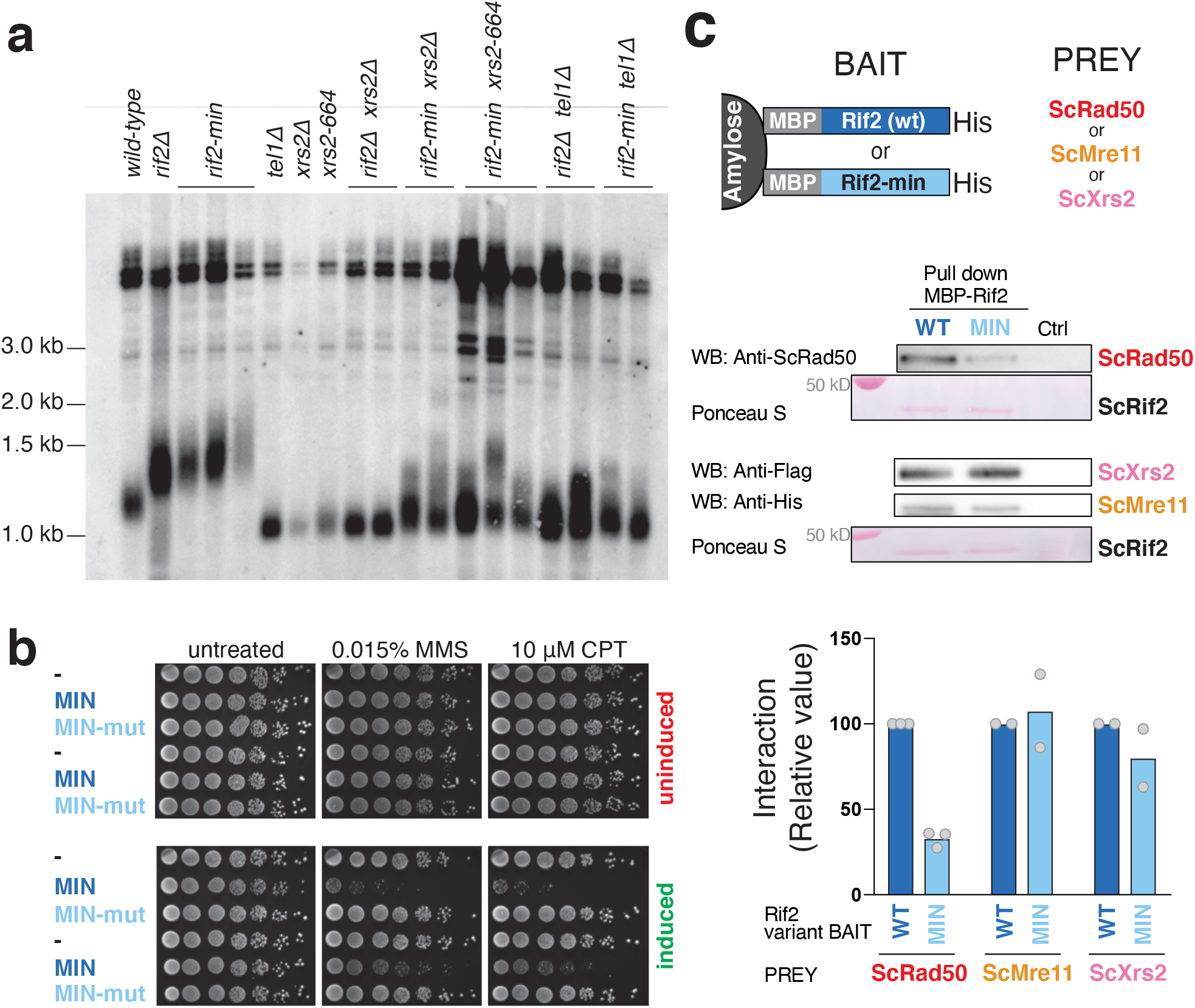
The MIN motif of Rif2 counteracts the action of the MRX complex in promoting telomere elongation and promotes binding to Rad50. A) Southern blot analysis of telomere length of *Saccharomyces cerevisiae* strains of the indicated genotypes. Genomic DNA was digested with XhoI and detected using a telomeric probe. ‘Δ’ indicates null alleles obtained by full deletion of the coding sequence. The *rif2-min* allele carries a F8E P10N substitution. B) Viability assays of strains overexpressing a fusion protein containing the first 34 amino acids of Rif2 under control of the galactose-inducible promoter, either in wild-type form (MIN) or mutant (MIN-mut, carrying the double F8E P10N substitution). Strains were grown in glucose-containing rich medium (YPAD) before plating onto either glucose (uninduced, top row) or induced (galactose, bottom row) plates containing the indicated chemicals and incubated at 30°C for 2-3 days. C) Protein interaction assays with immobilised wild-type or mutant Rif2 (Rif2-min) as bait, and Rad50, Mre11 or Xrs2 proteins used individually as prey. The bound proteins were separated by electrophoresis and detected by Western blotting (for Rad50, anti-Rad50; for Mre11-his, anti-his and for Xrs2-FLAG, anti-FLAG), or by Ponceau S staining (for Rif2 wild-type or Rif2-min). CTRL lane is prey protein incubated with amylose resin without bait. A representative blot is shown, as well as a quantification of 2-3 independent ones, as indicated.

If the MIN motif impairs the action of the MRX-Tel1 complex, it is possible that ectopic over-expression of only the N-terminal Rif2 region containing MIN, incapable to locate to telomeres due to the lack of binding to Rap1, might render cells hypersensitive to genotoxic stress, reminiscent of MRX mutants. In order to test this idea, we over-expressed peptides bearing the 34 amino acid residues of Rif2 from the strong galactose-inducible promoter. In line with the hypothesis above, we found that over-expression of the MIN motif, but not a mutant version bearing the F8E and P10N mutations, rendered the cells hypersensitive to treatment with methyl methanesulfonate (MMS) or camptothecin (CPT) (Figure 2b). Thus, expression of this large truncation allele of Rif2, containing only the MIN motif, has a dominant negative effect on the ability of cells to enforce DDR, consistent with a possible inactivating role on MRX.

Tel1 is pivotal in the telomerase-dependent pathway of telomere maintenance in budding yeast, but less crucial in assisting the DNA repair function of the MRX complex in budding yeast ^23,45^. In particular Tel1, is required to guarantee resistance to genotoxic stress by CPT but not MMS ^46^. If the MIN motif acted solely by affecting Tel1 recruitment to the complex, via the Xrs2 C-terminus, its overexpression might have been expected to have displayed specificity towards CPT sensitivity, instead of affecting sensitivity to MMS as well, as we observed (Figure 2b). We therefore decided to address whether Rif2 might function by targeting other subunits of the MRX complex instead of the Xrs2 C-terminus. In pull-down assays with purified components, Rif2 was found to be able to interact with all three subunits of the complex (Figure 2c). Interestingly, the F8E P10N mutations reduced, although did not abolish, the ability of Rif2 to interact with Rad50, but not with Mre11 and Xrs2, suggesting that the interaction with Rad50 might be the more relevant one with regard to the action of the MIN motif. Taken together, these results indicate that the MIN motif in Rif2 acts at telomeres by directly affecting the action of the MRX complex. In agreement with a recent biochemical study ^39^, this effect of MIN is likely exerted through an interaction on the Rad50 subunit. While that study has reported that an F8A mutation in Rif2, which impairs the ability of Rif2 to modulate MRX, does not affect the interaction with Rad50, the double mutant studied here shows diminished binding, suggesting that multiple contributions are made by MIN motif residues in contacting Rad50 ^39^.

### The MIN motif is required to execute the anti-checkpoint function of Rif2 at telomeres

Rif2 has a protective function at telomeres by both preventing exonucleolytic degradation and activation of the DNA damage response ^47,48^. Because the MRX-Tel1 complex contributes to the DDR response at unprotected telomeres, we tested the hypothesis that Rif2 might carry out its role in suppressing DDR activation at telomeres through the action of the MIN motif. To do so we employed a system first described by the Weinert lab that relies on the induction of a double-stranded break (DSB) by the HO endonuclease at a chromosomal site flanked by telomeric repeats ^49^. In our setting the DSB is flanked at both ends by telomeric repeats ^48^. The DNA break was induced, and small-budded cells, indicative of early S-phase stage, were micro-manipulated on a plate, and then inspected over several hours to measure the kinetics of progression through the cell cycle. Specifically, the length of the arrest in the G2/M phase following the break, as indicated by dumbbell-shaped cells, was monitored: while wild-type cells are expected to cycle quickly, as the DDR is not activated due to the protective effect of the telomeric repeats flanking the newly generated DNA ends, mutations affecting telomere protection will disable the anti-checkpoint function and result in transient cell cycle arrest in G2/M. In this assay, at least two independent pathways contribute to anti-checkpoint enforcement, one dependent on Rif1 and one dependent on Rif2 ^48^. We set out to test whether the contribution of Rif2 to checkpoint suppression relies on the MIN motif. As previously described, loss of Rif1 and Rif2 caused additive delays in release from G2/M in this assay (Figure 3a). Remarkably, in both the presence or absence of Rif1, the *rif2-min* allele produced the same effect as the *rif2* null allele (Figure 3a), indicating that the MIN motif is required for the ability of Rif2 to prevent checkpoint activation at unprotected telomeric DNA ends.

**Figure 3.**
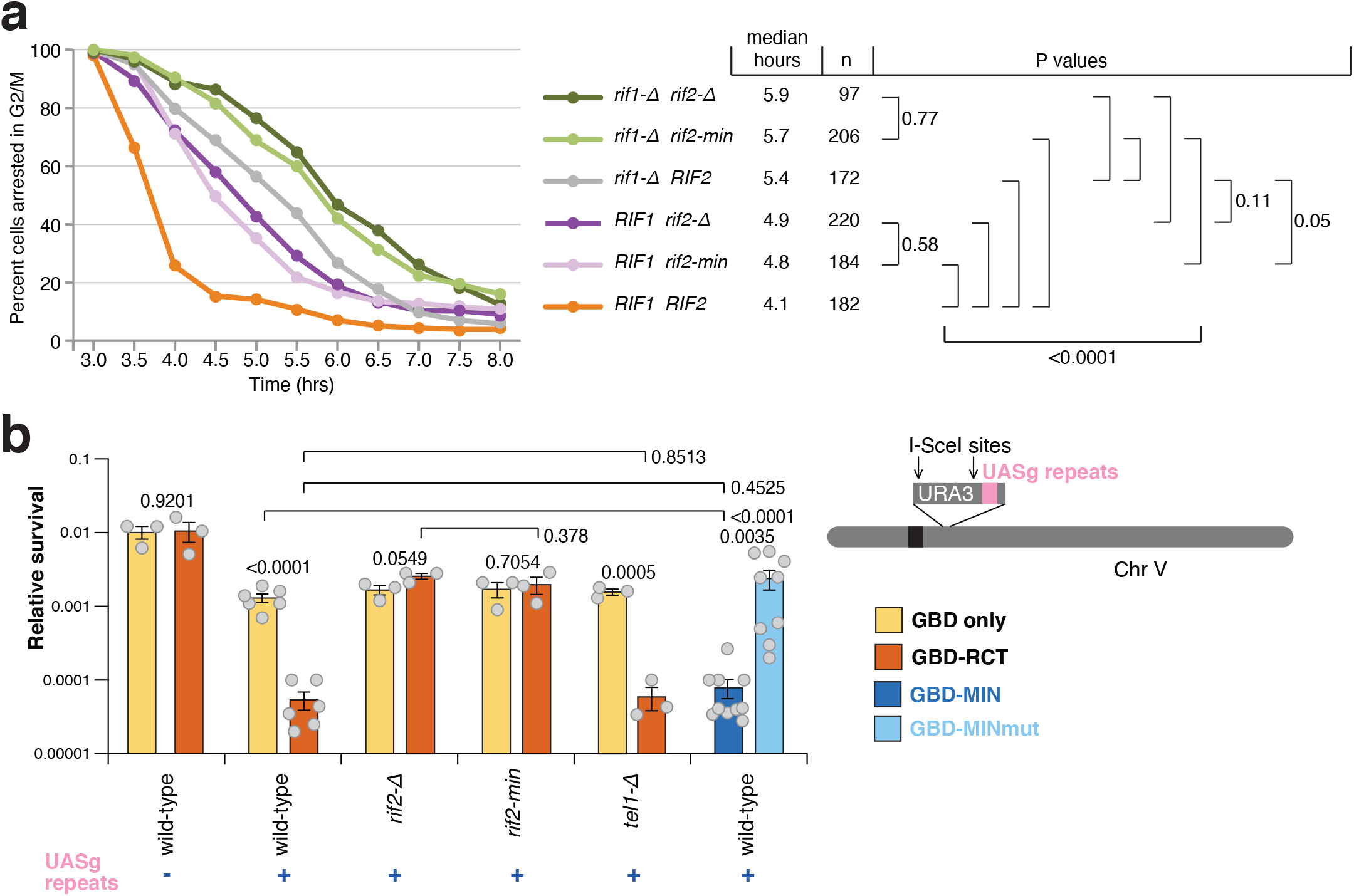
The MIN motif enforces suppression of the DDR and NHEJ at a DSB flanked by telomeric repeats. A) A specific DSB was induced at the ADH4 locus on the left arm of chromosome VII by expression of the HO endonuclease in galactose containing medium. After induction of the break, cells in S-phase were monitored for their ability to escape G2/M arrest. The percentage of G2/M arrested cells, from at least three independent experiments for each genotype (two for rif1-Δ), was estimated using Kaplan-Meier analysis by GraphPad Prism (Log-rank test). B) Viability assays to quantify the ability of cells to survive a double inducible DSB on either side of the URA3 gene on chromosome V. Strains were grown in minimal medium lacking uracil and tryptophan, then plated out on plates containing glucose (to assess plating efficiency) or galactose (to assess survival after cleavage). GBD is Gal4 DNA binding domain, RCT is the Rap1 C-terminal domain, MIN contains Rif2 amino acids 1-40, and the MINmut version carries the F8E P10N mutations. Relative survival was determined from at least three independent experiments. P values are reported for each pair of datasets as indicated, and were obtained with a two-tailed, unpaired t-test.

### The MIN motif of Rif2 suppresses NHEJ

In budding yeast, the MRX complex is required to promote DSB repair by NHEJ, and Rif2 suppresses NHEJ at telomeres ^33^. Because we showed that the Rif2 MIN motif limits the ability of MRX to promote telomere length, we addressed whether the motif might also be responsible for the disabling effect of Rif2 on NHEJ. DSB repair events can be channeled into the NHEJ pathway in the absence of an intact homologous sequence that could serve as template for repair by homologous recombination. NHEJ can be measured by assessing the ability of haploid budding yeast cells to survive a double DSB flanking a selectable marker ^33^. In this setting, telomeric sequences positioned near one of the two breaks cause decreased survival upon DSB induction, due to their suppression of NHEJ, at least partly via the Rap1-promoted recruitment of Rif2 ^33^. The effect of telomeric repeats in this assay can be bypassed by directly tethering the Rap1 C-terminal domain (RCT) near the DSB by fusing it to the Gal4 DNA Binding Domain (GBD) and placing copies of its recognition site (UASg) near the break. As expected, the presence of UASg sites near the break led to decreased survival upon the presence of a GBD-RCT fusion protein (Figure 3b). Deletion of Rif2 rescued the viability defect, due to loss of NHEJ inhibition and consequent DSB healing. Crucially the effect of the UASg/GBD-RCT combination depended on the presence of a functional MIN motif in Rif2, as the phenotypes of *rif2-Δ* and *rif2-min* strains were indistinguishable in this assay, indicating that the motif is required for the suppression of NHEJ by Rif2.

The observation that the role of the BAT domain in telomere length suppression did not require the RCT domain of Rap1 ^38^, predicts that tethering MIN directly to the break site, via the expression of a GBD-MIN fusion, should mimic the effect seen with the tethered RCT domain. Indeed, a decrease in survival was obtained when the first 40 amino acids of Rif2 were fused to GBD, while the F8E P10N double mutation abolished the NHEJ-suppressive effect of this small Rif2 N-terminal fragment fusion (Figure 3b). Taken together, these results indicate that the MIN motif of Rif2 is both necessary and sufficient to achieve the suppression of NHEJ in DSB repair by Rif2.

Interestingly, Tel1 was not required for the RCT to suppress NHEJ, suggesting that the effect of MIN in this setting occurs via the MRX complex itself, rather than via targeting the action of the Tel1 kinase, unlike in the case of the regulation of telomere length. This result is consistent with the suggestion, as described above, that the MIN motif acts via an interaction with Rad50, rather than by affecting the ability of Xrs2 to recruit Tel1, as originally proposed for Rif2. Because the *rif2-min* allele mimics the *rif2* null with regard to both telomere length and telomere protection we think it is unlikely that the MIN motif, and by extension Rif2, acts via the Xrs2-Tel1 recruitment axis, even in telomere length regulation. Rather these results are consistent with a model recently proposed whereby the Rif2 N-terminal region affects the ability of the MRX complex to activate the Tel1 kinase and specifically the ability of the Rad50 subunit to trigger its activation ^39^. Our results indicate that MIN affects a broad spectrum of MRX activities besides Tel1 activation, including specifically NHEJ.

### The MIN motif disables the endonucleolytic function of MRX

Our *in vivo* data indicate that the functions of Rif2 in telomere length regulation, suppression of the DNA damage checkpoint at telomeres, and inhibition of NHEJ at telomeres, are mediated by the MIN motif. In addition, epistasis analysis implicated the MIN motif of Rif2 in the control of MRX-Tel1, and mutating the motif diminished its ability to physically interact with Rad50, firmly linking the MIN motif with MRX functionality. To get insights into the mechanism of action of the MIN motif on the MRX complex, we expressed and purified recombinant full-length Rif2, in wild-type and the F8E P10N mutant form (Figure 4a), and tested its effect on various activities of the purified MRX complex. As observed previously ^23^, Rif2 was found to stimulate the ATPase activity of the MRX and MR complexes (Figure 4b). Importantly, the F8E P10N mutations ablated this stimulatory effect (Figure 4b). The ATPase activity of the Rad50 subunit is instrumental in promoting conformational changes in the complex, which modulate its activity in DNA end resection and tethering as well as Tel1 kinase activation ^20^. This raised the possibility that the MIN motif of Rif2 might directly impair the nucleolytic function of the MRX complex. To test this, we assayed nuclease activities of the MRX complex in the presence of wild-type and mutant Rif2. Previously, it has been established that MRX has a 3’-5’ exonuclease activity that does not require ATP hydrolysis (Figure 4c, left), and Mre11 alone is necessary and sufficient for this function ^50,51^. Additionally, MRX employs its endonucleolytic activity to resect DSBs with protein blocks. The endonucleolytic cleavage by the nuclease of Mre11 requires ATP hydrolysis by Rad50, as well as phosphorylated Sae2 (pSae2) as a co-factor (Figure 4c, right) ^28,52^. We employed a substrate used previously, which allows to monitor both exonuclease and endonuclease activities simultaneously (Figure 4c) ^52^. We observed that Rif2, whether in its wild-type or mutant form, had no effect on the exonuclease action of MRX, while wild-type Rif2, but not the Rif2-min mutant, was found to strongly reduce the endonuclease action of MRX in the presence of Sae2 (Figure 4d-i). This inability of the mutant to suppress endonucleolytic cleavage was not due to decreased DNA binding activity, as both mutant and wild type variants of Rif2 retained DNA binding capability (Figure 4j).

**Figure 4.**
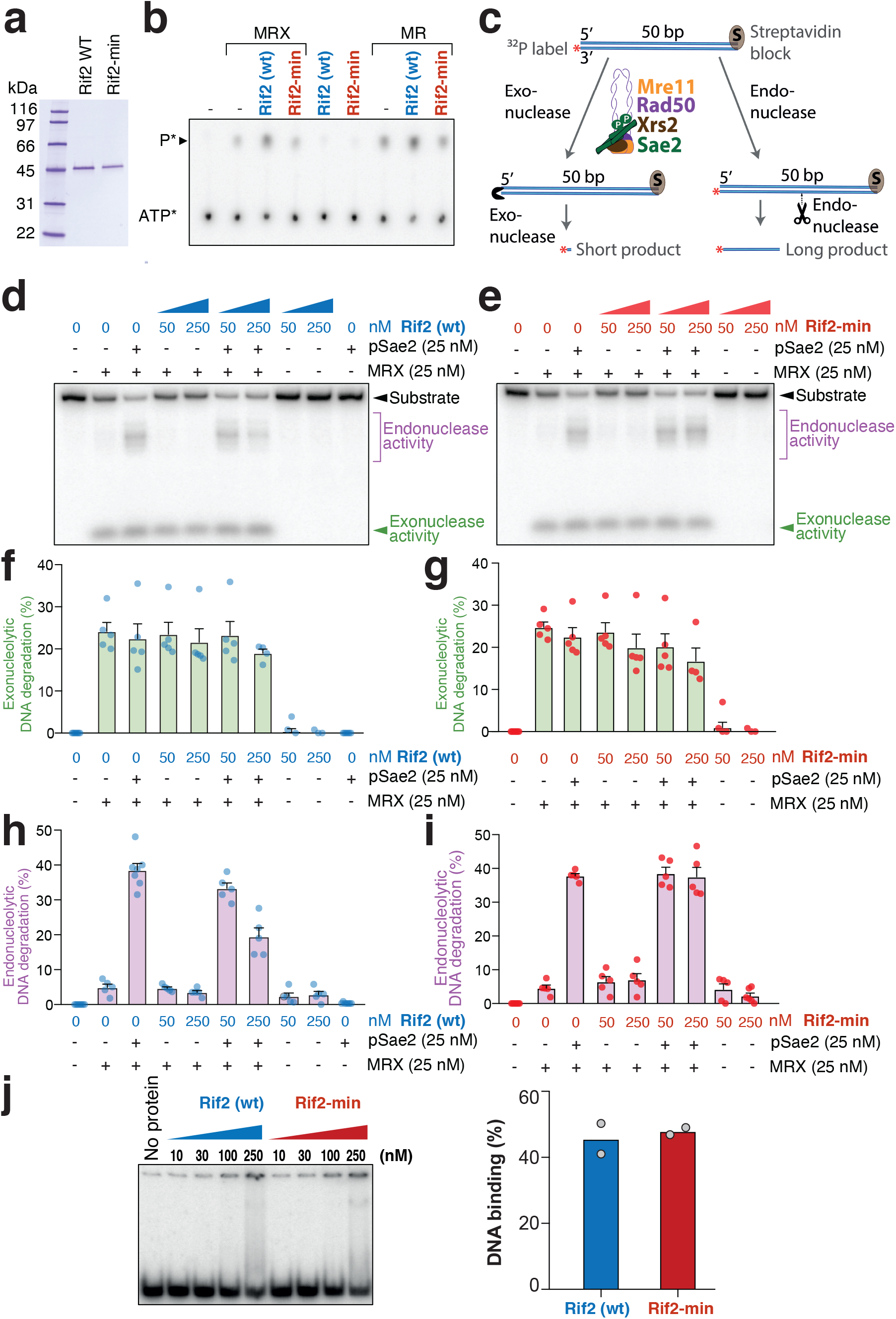
The MIN motif of Rif2 is required to suppress the endonucleolytic activity of MRX/MR. A) Purified *S. cerevisiae* wild type Rif2 and MIN-mutated Rif2-min variants used in this study. The polyacrylamide gel was stained with Coomassie Blue. B) Representative ATPase assay with MRX (100 nM), MR (100 nM), wild type Rif2 (500 nM) or Rif2-min (500 nM), as indicated. C) A scheme of the nuclease assay used to monitor the exonuclease and endonuclease activities of the MRX complex in the presence of phosphorylated Sae2. The radioactive label is at the 3’ position on the bottom DNA strand. The 3’-5’ exonuclease activity gives rise to a small product that migrates at the bottom of the gel. The endonuclease activity yields longer products. D) Nuclease assays with MRX, phosphorylated Sae2 (pSae2) and wild-type Rif2, as indicated. Shown is a representative experiment. E) Nuclease assays as in D, but with Rif2-min (F8E P10N mutations). F) and G) Quantitation of exonucleolytic data such as shown directly above in D) and E), respectively. Error bars represent SEM for 5 independent experiments. H) and I) Quantitation of endonucleolytic data as shown directly above in D) and E), respectively. Error bars represent SEM for 5 independent experiments. J) Left panel: representative electrophoretic mobility shift assays with Rif2 (wt) and Rif2-min with the indicated protein amounts. Right panel: quantitation of data such as in the gel shown at left, with 250 nM Rif2 (wt) or 250 nM Rif2-min; n=2, averages and individual data points are shown.

Rad50 is a key regulatory component in the MRX complex, which triggers ATP-dependent conformational switches to regulate its various functions. ATP binding by Rad50 is required for Tel1 activation by MRX and end tethering in NHEJ ^19,20,53^, and ATP hydrolysis is required for the Rad50-dependent endonucleolytic action of Mre11 ^28,50^. Accordingly, Rad50 mutants lacking ATP binding activity recapitulate the multiple phenotypes of the null allele ^54^. The ability of Rif2 to interact with Rad50, which is at least partly dependent on the MIN motif, and the requirement of the MIN motif to stimulate the Rad50 ATPase activity, suggest that Rif2 inactivates the multiple functions of MRX by triggering non-productive ATP hydrolysis by the MRX complex, and hence enforcing ATP-dependent conformational changes of the complex. It was recently shown that the F8 residue on Rif2 is instrumental in impairing the ability of Rad50 to activate Tel1, explaining the telomere length phenotype of the MIN motif mutants, which we here show is epistatic to the Xrs2-Tel1 pathway (Figure 2) ^39^. The suppression of NHEJ is also consistent with a MIN-dependent switch of the complex to an ‘open’ conformation after ATP hydrolysis. The nuclease data presented here further support this model, as we show that only the ATPase-dependent endonuclease activity of MR (or MRX) is affected, but not the ATPase-independent exonuclease activity. In this scenario, MIN is proposed to promote ATP hydrolysis by MRX (or MR) in a non-productive manner and thus prevent productive hydrolysis that would lead to endonucleolytic action. This conclusion agrees with previous data suggesting that Rif2 discharges the ATP-bound form of Rad50 to attenuates Tel1 activity ^39^. In summary, these results support a scenario where the MIN motif of Rif2 converts the MRX-ATP complex to the open ADP-bound form, which broadly impairs the MRX complex in its function to promote DNA repair, DDR signalling and telomerase regulation.

### The MIN motif of Rif2 interacts with the N-terminal region of Rad50

Among the characterized alleles of the MRX subunits, the *rad50S* allele stands out in that it leads to telomere elongation, rather than shortening as is seen in rad50 null cells ^10^. The *rad50S* alleles (generally represented by the Rad50-K81I mutation) comprise a small group of mutations originally identified for their meiotic phenotype in being defective in the resection of meiotic double strand breaks and removal of Spo11 from the breaks ^28,55^. The phenotype of rad50S mutants is similar in many respects to that of strains defective in the nuclease activity of Mre11 or lacking Sae2, which is required to stimulate the endonucleolytic action of Mre11 (reviewed in ^56^). *Rad50S* cells however differ in one significant aspect, telomere elongation, as *mreΔ* strains defective in endonucleolytic activity have normal telomere length, as do *sae2* null mutants ^57,58^. The failure to promote nucleolytic processing of DNA ends therefore cannot explain the elongated telomeres promoted by Rad50S.

The *rad50S* mutations are clustered in a small region at the N-terminus of the Rad50 protein, near the Walker A-type ATPase motif, in the part of the protein that contributes to half of the globular ‘head’ domain, and have recently been shown to define an area that is likely to provide the binding interface for phosphorylated Sae2, as the latter was shown to be uncapable of binding the Rad50-K81I protein ^52^. Prompted by the observation that the Rif2 MIN motif binds Rad50, we reasoned that the *rad50S* telomere phenotype could be explained if *rad50S* alleles failed to bind a repressor of MRX activity. We thus set out to test the idea that Rad50-K81I (from here onwards referred to as ‘Rad50S’) might be defective in MIN binding. To begin with, if the K81I mutation abolished the interaction with Rif2, then *rad50S* and *rif2-min* (or *rif2-Δ*) alleles would be predicted to be epistatic with regard to their effect on telomere length. As expected, single mutants showed telomere elongation, with *rif2* mutations conferring longer telomeres than *rad50S,* and *rif2-min* strains bearing slightly longer telomeres than *rif2* null ones (Figure 5a). Importantly, both *rad50S rif2-Δ* and *rad50S rif2-min* double mutant strains had telomeres of very similar length to *rif2* single mutants indicating that these *rad50* and *rif2* alleles are largely epistatic, and consistent with the idea that an interaction between the two proteins is compromised in both mutants.

**Figure 5.**
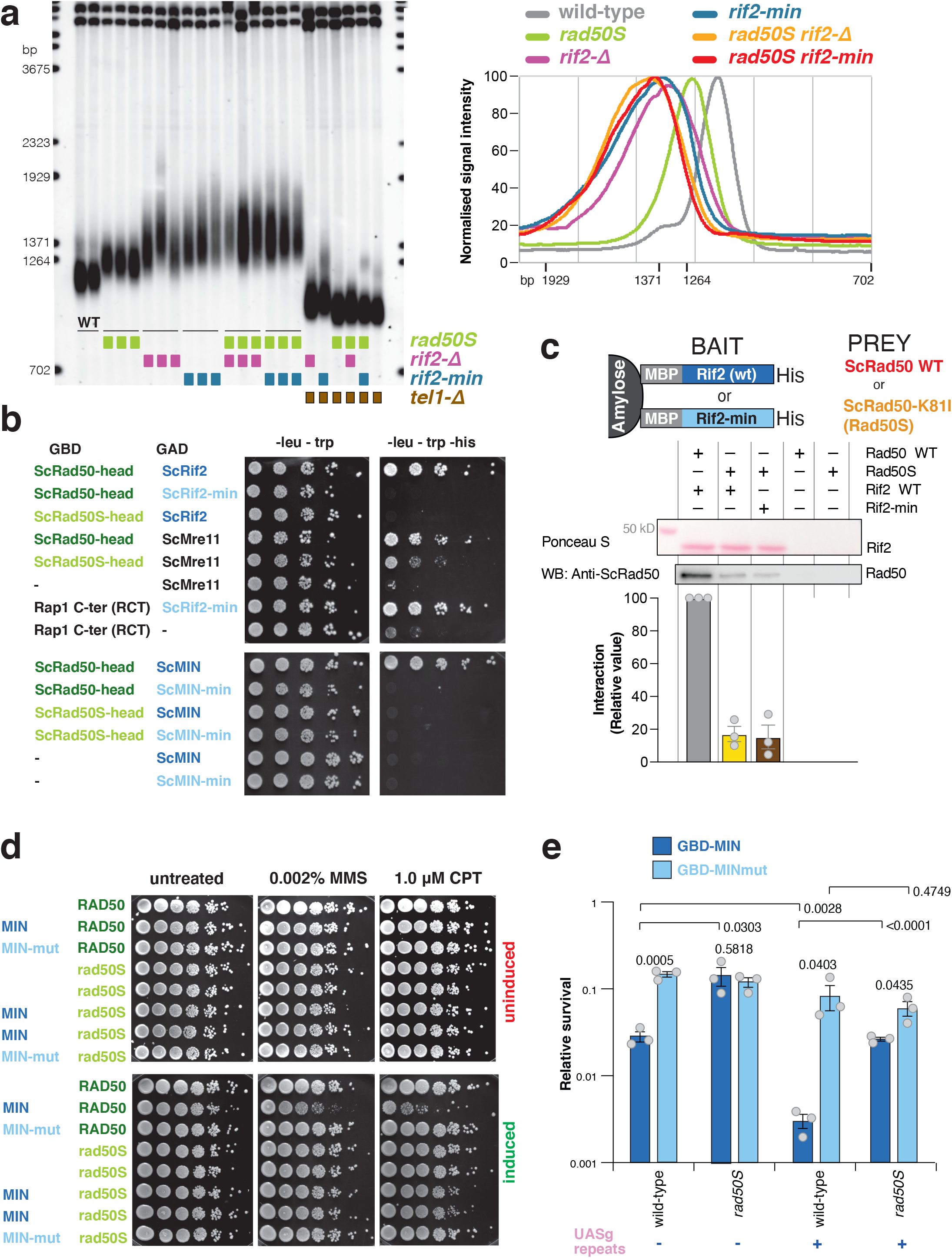
Rad50S (Rad50-K81I) fails to interact with the MIN motif of Rif2. a) Southern blot analysis of telomere length of *Saccharomyces cerevisiae* strains of the indicated genotypes, carried out as described in Figure 2a. A quantification of hybridization signals from this gel obtained with Image Gauge v4.1 (Fuji) is shown on the right panel; strains of identical genotypes were averaged together. b) Yeast two-hybrid assays of the *S. cerevisiae* fusion proteins indicated. Plates are minimal medium lacking the nutrients indicated on top. Five-fold serial dilutions were spotted left-to-right and incubated for 3-4 days at 30°C. c) Protein interaction assays with immobilized wild-type or mutant Rif2 (Rif2-min) as bait, and Rad50 or Rad50-K81I (Rad50S) proteins as prey. The bound proteins were separated by electrophoresis and detected by Western blotting (with anti-Rad50 antibody) or by Ponceau S staining (for Rif2 wild-type or Rif2-min). A representative blot is shown, as well as a quantification of 3 independent ones, as indicated. d) Viability assays of strains overexpressing a fusion protein containing the first 34 amino acids of Rif2 under control of the galactose-inducible promoter, either in wild-type form (MIN) or mutant (MIN-mut), as in Figure 2b. Uninduced strains were plated onto either glucose (uninduced, top row) or induced (galactose, bottom row) medium containing the indicated chemicals and incubated at 30°C for 2-3 days. e) Viability assays to quantify the ability of cells to survive a double inducible DSB on either side of the URA3 gene on chromosome V were carried out as in Figure 3b. GBD is Gal4 DNA binding domain, MIN contains Rif2 amino acids 1-40, and the MINmut version carries the F8E P10N mutations. Relative survival was determined for three independent strains; error bars represent SEM. P values are reported for each pair of datasets as indicated, and were obtained with a two-tailed, unpaired t-test.

To test this idea directly we performed yeast two-hybrid assays between wild-type and mutant variants of Rad50 and Rif2. Since we were interested to test interactions in the globular region of Rad50, we constructed a variant of the gene lacking the large central portion that codes for the coiled coils. As anticipated, we found that this variant, which we call Rad50head, is capable of interacting with Mre11 and, confirming the pull-down results with purified proteins (Figure 2c), also with full-length wild-type Rif2 (Figure 5b). Instead, the presence of either the *min* mutation in Rif2 (full-length and truncated) or the *S* mutation in Rad50, abolished the interaction in this assay. In agreement with this result we also found that mutant Rad50S was impaired in its interaction with Rif2 in the pull-down assay with purified proteins (Figure 5c). We note that the yeast two-hybrid assay appears to be more stringent as the interaction was not completely abolished in the pull-downs, no matter whether Rif2 or Rad50 was mutated (Figures 2c and 5c), leaving open the possibility that other regions of Rad50 might interact with Rif2 (multiple interactions for both Rif2 and Sae2 are observed with various components of MRX; Figure 2c ^52^). However, we did not detect additive effects when testing the two mutant variants in this assay (Figure 5c), suggesting that the specific interaction at this interface might be equally, and in terms of physiological relevance possibly completely, disrupted in the two mutants.

The interaction data support the epistasis analysis of telomere length in the mutants. To further evaluate the functionality of this interaction *in vivo* we decided to test the possible suppressive effect of the *rad50S* allele on two of the MIN phenotypes that we described earlier. The interpretation of the increased sensitivity of cells over-expressing MIN to genotoxic agents (Figure 2b) is that MIN directly interacts with MRX to disable its function in repair. So, if MIN requires the K81 residue to interact with Rad50, then the *rad50S* allele would be predicted to suppress the decreased viability for strains over-expressing MIN upon exposure to CPT and MMS. Indeed, in a *rad50S* background we observed a rescue of the MIN-dependent toxicity to both these drugs (Figure 5d). Similarly, we found that the ability of MIN to impair healing of a DSB by NHEJ when tethered near the break (Figure 3b) was partly compromised in strains bearing the *rad50S* allele (Figure 5e): in other words, *rad50S* suppressed the ability of MIN to inhibit NHEJ when tethered near the DSB. Interestingly, and in agreement with the MMS and CPT sensitivity data reported in Figure 2b, MIN was still able to inhibit healing of the break even in the absence of tethering, suggesting that overexpression of a functional MIN has an overall dampening effect on NHEJ within the cell. Taken together, these results identify the region in Rad50 that is responsible for a functional interaction with Rif2 via the MIN motif. Further studies will be required to elucidate the interplay between Rif2, Sae2 and Rad50 interactions but these data provide a molecular rationale for the longstanding observation of elongated telomeres in *rad50S* cells. The reason for the particular length setting in these cells, which is not as high as that seen in *rif2* mutants, is unclear. One possibility is that, although Tel1 activation after genotoxic stress in *rad50S* appears to be increased, the mutation might intrinsically impair full activation of Tel1 in the mutant complex even in the absence of Rif2 ^59^; alternatively the Rad50/Rif2 interaction might be more weakly affected *in vivo* by the rad50S mutation compared to the ones in MIN, although this is not supported by our biochemical data.

### A role for the MIN motif at telomeres beyond Rif2

Remarkably, as described above for the *Saccharomyces* genus (Figure 1a), all the yeast species analysed which bear both Rif2 and Orc4, display the motif in Rif2 but invariably lack it in Orc4 (Figure 6a) ^38,39^. These species belong to a clade within the Saccharomycetaceae that originated after the whole genome duplication event (species in red in Figure 6a). Instead, almost every species within the Saccharomycetaceae which lacked Rif2 displayed the MIN-containing N-terminal region Orc4 (Figure 6a, species in blue), with only two exceptions (species in black). A group of species within the post-WGD clade are devoid of Rif2, presumably because the duplicated *ORC4* gene was lost early, before it could differentiate into *RIF2*: these species also presented the MIN motif in Orc4. The analysis suggests that the MIN motif originated in Orc4 and that its function was then retained in Rif2 and lost from Orc4 (presumably because redundant) in the budding yeast *Saccharomyces cerevisiae* and closely related species.

**Figure 6.**
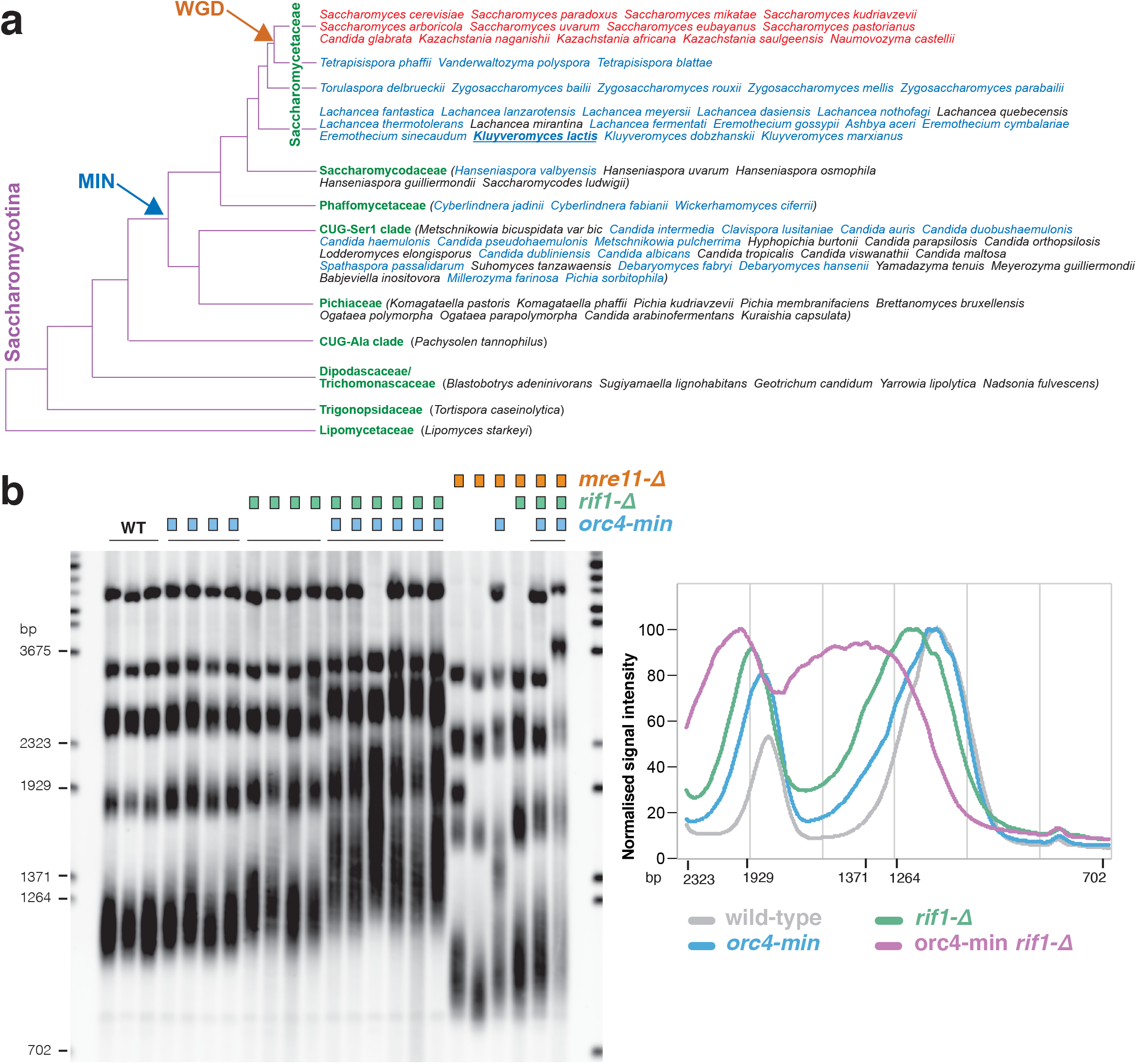
The MIN motif of *K. lactis* Orc4 suppresses telomere elongation. a) Phylogenetic tree of the Saccharomycotina, adapted from ^73^. Only species for which a search for Rif2 and/Orc4 was carried out are included. Species bearing Rif2 are in red; in all cases Rif2 has MIN domain whereas Orc4 does not. Species bearing putative MIN domain in Orc4 are in blue, whereas those where MIN domain is not present in Orc4 are in black. All red- and blue-lettered species have their MIN sequenced reported in Figure 1B. The occurrence of the Whole Genome Duplication (WGD) event during the evolution of the Saccharomycetaceae clade is indicated by the orange arrow, whereas the deduced appearance of the MIN motif in Orc4 within the Saccharomycotina is indicated by the blue arrow. *K. lactis* is underlined. b) Southern blotting of EcoRI-digested DNA hybridised with a telomeric oligonucleotide for the indicated genotypes. A quantification of hybridization signals from this gel obtained with Image Gauge v4.1 (Fuji) is shown on the right panel; strains of identical genotypes were averaged together.

The proposition that the MIN motif has originated in Orc4 coupled with the suggestion that MIN in Rif2 acts to regulate the action of MRX-Tel1 in budding yeast, raised the possibility that Orc4 might have originally had a similar role in affecting MRX-Tel1 before this function was taken up by Rif2. In order to test this idea, we turned to *K. lactis*, since this yeast (underlined in Figure 6a) does not bear Rif2 and has a MIN-carrying Orc4. Similarly to budding yeast, MRX in *K. lactis* is required to maintain telomere length ^60^, and mutations in the MIN motif of KlOrc4 would thus be predicted to lead to telomere elongation. We introduced the F8E and P10N mutations into KlOrc4 (*orc4-min*) and measured telomere length in the resulting strains. Although some variation was observed, a moderate level of telomere elongation was observed in *orc4-min* isolates, as longer telomeres were over-represented in these strains compared to wild-type (Figure 6b). Because the Kl *orc4-min* and Sc *rif2-Δ* telomere length phenotypes are relatively minor in comparison to that enforced by the lack of Rif1 in either species (Figure 6b) ^32^, to further confirm the elongation phenotype of the *orc4-min* allele, we combined it with a null mutation in *rif1*, since in budding yeast disabling both the Rif1- and Rif2-dependent pathways of telomere elongation has a synergistic effect ^32^. Indeed, combining the *rif1-Δ* and *orc4-min* mutations led to a clear additive effect on telomere elongation, indicating that Rif1 and Orc4 independently regulate telomere length in *K. lactis*, the latter protein doing so through its MIN motif. We next combined the *orc4* and *rif1* mutations with a *mreΔ* deletion allele, (Figure 6b). Telomeres in the double and triple mutants were drastically shortened, indicating that *MREΔ* is largely epistatic to *ORC4* and *RIF1* in telomere length regulation in *K. lactis*, consistent with the idea that Orc4 MIN acts by affecting MRX-Tel1. These results suggest an unanticipated role for Orc4 at telomeres and are consistent with subfunctionalisation of the Orc4 and Rif2 proteins after the WGD event, where Orc4 has retained a function in DNA replication while Rif2 has taken up the role in MRX-Tel1 control at telomeres. It remains to be established whether Orc4 has a role in *K. lactis* telomeres as part of the ORC complex or whether it acts independently of it. As Rif2 is delivered to telomeres via the RCT of Rap1, one possibility is that Orc4 might also be recruited via an interaction with Rap1. Residues that are important in Rif2 for interaction with the RCT ^61^ do not seem to be particularly well conserved in Orc4, and a bias towards conservation in the MIN-bearing Orc4 proteins is not apparent. Furthermore, some species that have an Orc4 MIN do not have a Rap1 RCT (for example *Candida albicans*). It remains therefore unclear whether the RCT of Rap1 might play a role in recruiting Orc4 at telomeres in some settings ^62^, but we note that the role of human telomeric protein TRF2 in recruiting ORC to telomeres has been reported ^63^. Both the dynamics and mechanism of recruitment of Orc4 to fungal species will require further investigation.

In seeking to understand how general a role MIN might play in regulation of the MRX complex, we sought to determine how widespread MIN is in ORC4 genes. Within the Saccharomycotina, the MIN motif was found to be present in the Phaffomycetaceae and CUG-Ser-1 clades, and possibly in one species within the Saccharomycodaceae (Figure 6a, species in blue). In the Ascomycota, we failed to identify the MIN motif in any of the remaining groups within the Saccharomycotina (see four bottom clades in Figure 6a), as well as in the two other groups which together with the Saccharomycotina comprise this group (the Pezizomycotina - 4 species examined - and the Taphrinomycotina - 12 species); we also could not find it in the Basidiomycota (3 species) (see Figure 1e). It therefore appears likely that the MIN motif evolved in Orc4 at some point within the Saccharomycotina, and that several species within this group lost it in Orc4 afterwards. Although we conducted searches for the motif in the genomes of some of the candidates where it would be most strongly predicted to be present, due to their belonging to clades where it is prevalent (for example *Lachancea quebecensis* and *mirantina*, and *Candida tropicalis*), we failed to find clear hits. It is unclear how the function of MIN is fulfilled at the telomeres of species that are devoid of it, but these results are consistent with the emerging remarkable modularity and flexibility of the telomeric complex in dealing with its multiple tasks (within shelterin, for example, several differences have been documented between the human and mouse complex, reviewed in ^7^).

The fission yeast *Schizosaccharomyces pombe* is a well-studied model system for telomere biology. When we searched for the MIN motif in *S. pombe* we found a match in Taz1. In *S. pombe* the telomeric complex does not rely on Rap1 for binding to telomeric DNA repeats, but utilizes instead the DNA binding protein Taz1. Remarkably, this putative MIN motif was conserved in all four *Schizosaccharomyces* species whose genome has been sequenced (Figure 7a). Although several residues from the extended consensus derived from Rif2 and Orc4 sequences, particularly in its N-terminal region, were absent, the four invariant residues in the core motif at positions +1, +2, +5 and +6 (Figure 1c) were present. The highly conserved proline at +4 was also present, and a stringent requirement seems to operate in *Schizosaccharomyces* at the otherwise relaxed position +3. Intriguingly, the motif is located in the C-terminal part of the protein, in a small region of little sequence conservation outside the MIN motif itself and of no known function, which is sandwiched between the Myb/SANT DNA binding domain and a homodimerization domain (Figure 7b). In *S. pombe* strains carrying the same F to E and P to N mutations (*taz1-min*) equivalent to those characterised in *S. cerevisiae*, we did not observe strong changes in telomere length: this is not surprising as telomere length is not dependent on MRN-Tel1 in this species ^64^. Similarly we failed to observe strong defects in the telomere protection in this mutant. In fission yeast, deprotected telomeres undergo fusions in a manner dependent on MRN when cells are kept in a prolonged G1 arrest. Strains carrying *taz1-min* showed no obvious effect on cell viability (data not shown). This result is consistent with the major role played by Rap1 in the suppression of NHEJ in *S. pombe* ^65^.

**Figure 7.**
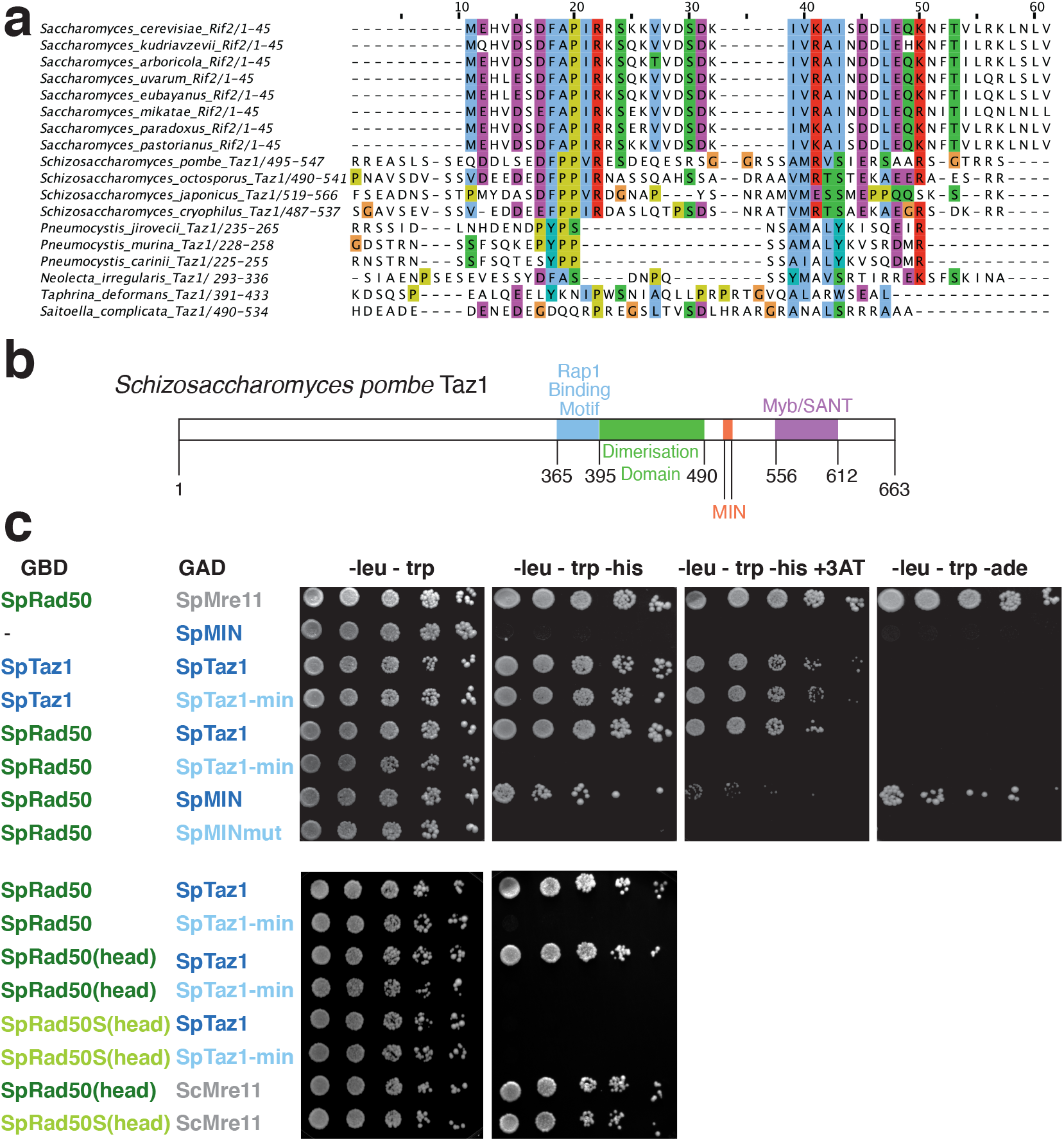
A MIN motif in fission yeast Taz1 binds the Rad50 N-terminus. a) Sequences of putative Taz1 proteins from the Taphrinomycotina were aligned using Clustal Omega with the N-terminal region of Rif2 from Saccharomyces species. b) Diagram of the domain structure of *Schizosaccharomyces pombe* Taz1. Domains previously identified in SpTaz1 ^75^ are indicated for the C-terminal half of the protein. The MIN motif is in orange. c) Yeast two-hybrid assays of the *S. pombe* fusion proteins indicated. Plates are minimal medium lacking the nutrients indicated on top (and one instance containing 3 mM 3-aminotriazole). Five-fold serial dilutions were spotted left-to-right and incubated for 3-4 days at 30°C.

Because telomeres possess redundant pathways to disarm the various branches of the DNA damage response, we hypothesise that the MIN motif in *S. pombe* might act redundantly with other telomere factors in dealing with the threat posed by untimely activation of MRN. We decided to address this possibility by assessing the ability of the Taz1 MIN to interact with MRN, by using yeast two-hybrid assays. Whereas no interaction between Taz1 and either Mre11 or Nbs1 was detected with this assay (data not shown), an interaction between Taz1 and Rad50 was readily observed (Figure 7c). Crucially, the interaction was abolished when the Taz1-min (F511E, P513N) protein was used, while this protein retained, as expected, the ability to homodimerize, and is therefore functional. Finally, binding to Rad50 was retained when only the region of Taz1 between the dimerization and DNA binding domains was included (amino acids 489-558); again, introduction of the F511E, P513N double mutation abolished the ability of the 70 amino acids spanning the MIN motif to interact with Rad50. These results show that the MIN motif is required for the interaction of Taz1 with Rad50 and that a small part of the Taz1 protein overlapping the MIN motif is sufficient for this interaction, and thus mimic and extend the interaction data obtained with Rif2 and Rad50 in budding yeast. Strikingly, a K81I mutation in *S. pombe* Rad50, equivalent to the same change in the *S. cerevisiae* protein (Rad50S) abolished the interaction between Rad50 and Taz1, strongly suggesting a conserved mode of interaction between Rif2/Taz1 and Rad50 (Figure 7c, bottom panels). These results predict that Taz1 interacts with Rad50 in fission yeast cells. It remains to be seen whether the Taz1 motif has a similar role to Rif2 in modulating the activity of MRN. Because Taz1 is evolutionarily unrelated to Rif2 or Orc4, we suggest that the MIN motif has appeared independently in the Schizosaccharomycetes. We also note the motif is not present in other Taz1 proteins from the Taphrinomycotina (Figure 7a), indicating that the motif appeared in this clade after the appearance of Taz1, which is of uncertain origin but might be linked to the transcription factor Tbf1.

## Conclusion

We presented evidence that several yeast species have evolved a protein motif that has the remarkable ability to dismantle the multiple functionalities of the MRN/X complex at telomeres. The available evidence suggests that the MIN motif achieves this by promoting non-productive ATP hydrolysis by the MRN/X complex, and thus abolishing the functions of MRN/X that depend on productive ATP hydrolysis. Our biochemical data suggest that the MIN motif appears to hijack a regulatory step normally leading to the activation of the endonuclease activity of MRN/X to instead disable it. In budding yeast, the action of MRX-Tel1 is not only a potentially dangerous trigger for DDR activation but is also required for telomerase action, which occurs preferentially at shorter telomeres. Mre11 appears to facilitate telomerase recruitment specifically to the leading strand telomere, possibly by promoting its resection ^66^. Remarkably, Rif2 marks the shorter telomeres for preferential telomerase action ^67^, so it is possible that maintaining a threshold of telomere-bound MIN motifs is required for efficient MRN inhibition. This mechanism could potentially help limit spurious action of MIN at non-telomeric sites, where its nucleation might not reach the required threshold ^23^. Similar considerations might apply to fission yeast telomeres, but the exact role for the Taz1 MIN motif remains to be determined. We note that human cells actively recruit MRN to telomeres ^68,69^, and that a role of ATM in telomerase activation has recently been documented ^12,13^. In addition, MRN can have a protective role at mammalian telomeres, by preventing telomere fusions ^70,71^. Thus it is possible that the fine tuning of MRN action in higher Eukaryotes requires strategies similar to those allowed by the deployment of the MIN motif in yeast. It remains to be seen how the MIN motif might affect the N-terminal region of Rad50, and how its lodging at this location might trigger conformational changes in the rest of the molecule ^39^. Interestingly, *rad50S* alleles were found to lead to increased telomere fusions and genome instability in mice, raising the possibility that similar mechanisms to control MRN action at telomeres might exist in higher Eukaryotes ^72^.

## Methods

### Strains and plasmids

All *Saccharomyces cerevisiae* (*Sc*) strains used in this study were derivatives of W303 (*leu2-3,112 trp1-1 can1-100 ura3-1 ade2-1 his3-11,15). Sc* and *Kluyveromyces lactis* (*Kl*) strains were grown in YPAD rich medium or synthetic complete (SC) minimal medium (2% w/v glucose, 0.67% w/v Yeast Nitrogen Base, 0.2% w/v SC dropout mix, 2% w/v agar, pH 6.0). YNB and SC mixes were from United States Biological. For transformations, the LiAc protocol was followed, with heat shock at 42°C for 45 min.

The *rif2-min* allele was generated by transformation of *Sc* strains with plasmid pAB1703, which contains a small genomic *RIF2* regions into which the F8E P10N mutations were engineered: the plasmids was linearised with BsrGI before transformation, transformants were selected on SC-ura medium and then counter-selected on 5FOA-containing medium to identify recombination events that popped-out the inserted plasmid from the *RIF2* locus. Transformants that retained the mutations were screened by PCR using the silent BglII introduced alongside the mutations. A similar approach was used for the *orc4-min* allele in *Kl* using plasmid pAB2049 linearised with EcoRI.

For over-expression of the Rif2 MIN motif in *Sc*, two plasmids were used, containing the first 34 amino acids of Rif2, either in wild-type or F8E P10N form, N-terminally fused to a GFP Binding Protein fragment, expressed from a galactose-inducible promoter (pAB2081 and pAB2082, respectively). The plasmids were linearised with BstXI before before being transformed into *Sc* strains. Integration of the plasmids at the *HIS3* locus was confirmed by PCR.

The *rad50S* allele was introduced in the required genetic background using pNKY349 (pAB2301) cut with Cut with BamHI-EcoRI. Correct transformants were screened by PCR and MMS sensitivity. Transformants that bore the correct *URA3* integration but retained the wild-type *RAD50* allele after recombination of the construct, were also selected and used in the CPT and MMS sensitivity experiments to ensure equal assortment of auxotrophic markers in spotting assays. For the NHEJ experiments, a variant of the pNKY349 plasmid where the URA4 marker was swapped with HIS3 was constructed, pScRad50-K81I-Hi (pAB2304) and used in the same way.

### Southern blotting analysis

<u>*Sc*</u> or *Kl* genomic DNA was prepared from 10 ml log-phase cultures in YPAD rich media by vortexing cell pellets with 0.3 g glass beads in 0.4 ml 2% Triton-X, 1% SDS, 100 mM NaCl, 10 mM Tris-HCl pH8.0, 1 mM EDTA pH8.0, followed by phenol:chloroform:isoamyl alcohol (25:24:1) extraction, ethanol precipitation and RNase A digestion. For telomere length analysis, 1 µg of genomic DNA was digested with XhoI (*Sc*) or Eco RI (*Kl*) overnight and gel electrophoresis was carried out on 1% agarose gels. Depurination was carried out by soaking gel in 125 mM HCl for 10 min, followed by denaturation in 0.5 M NaOH 1.5 M NaCl for 30 min. Overnight transfer was in 20x SSC using nylon membranes (Roche). After UV crosslinking, membranes were incubated at 60°C (Sc) or 50°C (Kl) in 5x SSC, 5% dextran sulphate, 0.2% I-Block (ThermoFisher T2015), 0.1% SDS. DNA probes were briefly denatured before being added, and hybridisation was carried out overnight. Washes were at room-temperature with buffers pre-heated at 60°C (Sc) or 50°C (Kl) twice in 1x SSC 0.1% SDS then twice in 0.5x SSC 0.1% SDS. Detection was by incubation for 1 hr at RT in 100 mM Tris pH7.55, 150 mM NaCl, 1% skimmed milk, plus 1µI of alkaline phosphatase tagged anti-fluorescein F(ab) (Roche), followed by four washes at RT in 100 mM Tris pH7.55, 150 mM NaCl, and detection with CDP-Star detection agent. Imaging was on a LAS4000 instrument. Probes were made either by PCR of a Y’ fragment followed by random priming (*Sc*) or by terminal transferase extension of a 25-mer telomeric oligonucleotide (*Kl*), both in the presence of Fl-dUTP.

### Single-cell checkpoint arrest analysis

Strains were grown overnight in 5 ml of YPLG (acid lactic and glycerol) medium. The following morning, cells were diluted into 5 ml of the same medium to 3*10^5^ cells/ml and incubated at 30°C. After 2 hours of incubation, 1 ml of 20% galactose was added in order to induce the HO endonuclease gene and the strains were incubated for further 2 hours. After that, the cells were plated down in the central line of a YPAD plate, kept previously at room temperature. Then small budded cells were dissected into a grid by a micromanipulator for analysis. After 1 hour from the beginning of the dissection, cells were checked every 30 minutes for the second round budding in order to record the cell-cycle restart. YPAD plates were preserved at 30°C into an incubator between observations. After growth of the colonies at 30°C, the following day, the central line of the plate was removed and it was incubated for further one day. The next morning the strains were replicated in Sc-Lys plate and incubated for two days at 30°C. In the end, Lys^**+**^ colonies were excluded from the dataset because the LYS2 gene is located on the telomere-proximal side of the HO site and only cells Lys^**−**^ that had been subjected to a DNA break were scored. Average restart time for each construct was estimated using Kaplan-Meier survival analysis in GraphPad.

### NHEJ assays

Strains were grown on solid minimal medium containing glucose and lacking uracil and tryptophan, then grown to saturation in the same medium overnight, before being diluted and plated on the same plates (to calculate plating efficiency, used for normalisation) and also on minimal plates lacking tryptophan and containing glucose ^33^.

### Yeast two-hybrid analysis

Yeast two hybrid assays were performed by co-transforming GBD (Gal4 DNA Binding Domain) and GAD (Gal4 Activating Domain) plasmids (in the pGBKT7 and pGADT7 verctors) in various combinations in the PJ69-4A budding yeast strain (*MATa trp1-901 leu2-3,112 ura3-52 his3-200 gal4 gal80 LYS2::GAL1-HIS3 GAL2-ADE2 met2::GAL7-lacZ* ). Transformants were screened for interaction by spotting 5-fold dilutions on minimal plates lacking tryptophan and leucine for selection of GBD and GAD plasmids respectively. Interactions were assessed by quantification of expression of the *HIS3* and *ADE2* markers by plating on minimal medium lacking, in addition, histidine or adenine. The stringency of the *HIS3* reporter was also increased by the addition of 3 mM 3-aminotriazole. Plated were incubated at 30°C for 3 (SC-leu-trp) or 4 (selective plates) days.

### Recombinant protein purification

All proteins were expressed in *Spodoptera frugiperda* 9 (*Sf*9) cells according to standard procedures. *S. cerevisiae* Mre11-Rad50-Xrs2 (MRX, containing Mre11-his, Xrs2-FLAG and untagged Rad50), Mre11-Rad50 (MR, containing Mre11-his and untagged Rad50), Mre11 (his-tagged at C-terminus), Rad50 (FLAG-tagged at C-terminus), Rad50S (K81I, FLAG-tagged at C-terminus) and Xrs2 (FLAG-tagged at C-terminus) proteins were prepared as described previously ^28,51,52,76^. Phosphorylated Sae2 (pSae2, his-tagged as C-terminus) was prepared as described previously ^52^.

Wild type Rif2 and Rif2-min variants were expressed using pFB-MBP-Rif2-his or pFB-MBP-Rif2-min-his vectors, respectively. The *Sf*9 cell pellets (from 1000 ml culture) were resuspended in 4 volumes lysis buffer (50 mM Tris-HCl pH 7.5, 1 mM dithiothreitol, 1 mM ethylenediaminetetraacetic acid, 1:400 Sigma protease inhibitory cocktail P8340, 1 mM phenylmethylsulfonyl fluoride, 30 μg/ml leupeptin) and incubated for 20 min at 4°C. 1/2 volume 50% glycerol and 6.5% volume 5 M NaCl solutions were then added, and the sample was incubated for 30 min while stirring. The cell extract was centrifuged (48,000 g for 30 min) to obtain soluble extract. The extract was bound to 6 ml amylose resin (New England Biolabs) for 1 h batchwise. The resin was washed with wash buffer (50 mM Tris-HCl pH 7.5, 5 mM 2-mercaptoethanol, 1 M NaCl, 10% glycerol, 1 mM phenylmethylsulfonyl fluoride), and protein was eluted with elution buffer (50 mM Tris-HCl pH 7.5, 5 mM 2-mercaptoethanol, 300 mM NaCl, 10% glycerol, 1 mM phenylmethylsulfonyl fluoride, 10 mM maltose). The protein concentration was estimated by the Bardford method and PreScission protease (1:5, w/w) was added, and the sample was incubated for 2 h at 4 °C. Then, 10 mM imidazole (final concentration) was added to the sample and the solution was bound to 0.5 ml NiNTA resin (Qiagen). The resin was washed with NTA buffer A1 (50 mM Tris-HCl pH 7.5, 5 mM 2-mercaptoethanol, 1 M NaCl, 10% glycerol, 0.5 mM phenylmethylsulfonyl fluoride, 58 mM imidazole) and NTA wash buffer A2 (50 mM Tris-HCl pH 7.5, 5 mM 2-mercaptoethanol, 150 mM NaCl, 10% glycerol, 58 mM imidazole) and eluted with NTA buffer B (50 mM Tris-HCl pH 7.5, 5 mM 2-mercaptoethanol, 150 mM NaCl, 10% glycerol, 300 mM imidazole). The eluate was dialysed into 50 mM Tris-HCl pH 7.5, 5 mM 2-mercaptoethanol, 150 mM NaCl, 10% glycerol, aliquoted, frozen in liquid nitrogen and stored at −80 °C. The purification yielded approx. 1 ml of 10 μM Rif2 (his-tagged at C-terminus). The Rif2-min variant was prepared in the same way.

### ATPase assays

The reaction buffer contained 25 mM Tris-acetate pH 7.5, 5 mM magnesium acetate, 1 mM dithiothreitol, 0.25 mg/ml BSA (New England Biolabs), 150 μM cold ATP, 1 nM labeled y-^32^P ATP, 50 nM DNA substrate (in molecules, annealed oligonucleotides 210- GTAAGTGCCGCGGTGCGGGTGCCAGGGCGTGCCCTTGGGCTCCCCGGGCGCGTACTCCACCTCATGCATC and 211- GATGCATGAGGTGGAGTACGCGCCCGGGGAGCCCAAGGGCACGCCCTGGCACCCGCACCGCGGCACTTAC) and respective recombinant proteins. The reactions were incubated for 4 h at 30 C. The reactions were stopped with EDTA (50 mM final concentration). The substrate and hydrolysed ATP were separated by thin layer chromatography using 0.3 M lithium chloride and 0.3 M formic acid (mixed 1:1) as the mobile phase. The plates were then dried and exposed to storage phosphor screens and signal was detected using Typhoon imager (GE Healthcare).

### Nuclease assays

Nuclease assays with purified proteins were carried out as described previously ^52^. The 50 bp-long DNA substrate was prepared by annealing oligonucleotides PC1253C (AACGTCATAGACGATTACATTGCTAGGACATCTTTGCCCACGTTGACCCA) and PC1253B (TGGGTCAACGTGGGCAAAGATGTCCTAGCAATGTAATCGTCTATGACGTT) with 3’-terminal biotin ^28^. Upon annealing and purification, the substrate was bound to streptavidin to create a protein block. Briefly, the nuclease buffer contained 25 mM Tris-acetate pH 7.5, 1 mM dithiothreitol, 0.25 mg/ml BSA (New England Biolabs), 1 nM DNA substrate (in molecules), 5 mM magnesium acetate, 1 mM manganese acetate, 1 mM ATP, 80 units/ml pyruvate kinase, 1 mM phosphoenolpyruvate and the recombinant proteins, as indicated.

### Electrophoretic mobility shift assays

To monitor DNA binding, assays (15 μl) were carried out in 25 mM Tris-acetate pH 7.5, 1 mM dithiothreitol, 5 mM magnesium acetate, 1 mM manganese acetate, 0.25 mg/ml BSA (New England Biolabs), 1 mM phosphoenolpyruvate, 1 mM ATP, 80 U/ml pyruvate kinase, and 1 nM ^32^P-labeled dsDNA substrate (in molecules, annealed oligonucleotides PC1253C and PC1253B). Afterwards, recombinant proteins were added, and the reactions were incubated at 30 °C for 30 min. Afterwards, the samples were mixed with 4 μl 50% glycerol with bromophenol blue, and separated by electrophoresis in 6% polyacrylamide in TAE buffer. Gels were dried and developed by autoradiography.

### Pulldown assays

The *Sf*9 cell lysate containing approx. 5 μg of MBP-Rif2 or MBP-Rif2-min (see recombinant protein preparation) was incubated for 1 h at 4 °C with amylose resin (New England Biolabs, 50 μl per reaction). The resin with the immobilized MBP-Rif2 variants was washed 5 times with 1 ml wash buffer (50 mM Tris-HCl pH 7.5, 200 mM NaCl, 0.2% NP40, 2 mM EDTA, Sigma protease inhibitory cocktail P8340 1:1000), using 2000 rpm for 2 min centrifugation each time to pellet the resin. Afterwards, 1 μg of Mre11, 500 ng of Rad50, or 1 μg of Xrs2 were added to the resin and incubated together for 1 h in the same wash buffer. Afterwards, the resin was washed 4 times with 1 ml wash buffer to remove unbound proteins. Bound proteins were then eluted with 100 μl wash buffer supplemented with 20 mM maltose. Prescission protease was then added (0.5 μg) to cleave MBP tag from MBP-Rif2 or MBP-Rif2-min. The proteins in the eluate were separated by electrophoresis and detected by Western blotting. The antibodies used were: anti yRad50, Thermo Scientific, PA5-32176, 1:1000; anti FLAG, Sigma, F3165, 1:1000; anti-His, MBL, D291-3, 1:5000. Pulldown assays with the Rad50 K81I variant were carried out as above with the following modifications: the resin with the immobilized MBP-Rif2 variants was first washed 4 times with 1 ml wash buffer containing 1M NaCl (50 mM Tris-HCl pH 7.5, 1 M NaCl, 0.2% NP40, 2 mM EDTA, Sigma protease inhibitory cocktail P8340 1:1000), then once with the same buffer but containing 150 mM NaCl. Afterwards, 750 ng of recombinant Rad50 or Rad50K81I were added to the resin and incubated for 1h at 4 °C in wash buffer containing 150 mM NaCl. The same buffer was then used in washes to remove the unbound proteins, as described above.

## Acknowledgements

We thank Lorraine Symington, David Shore, Jürgen Heinisch, Yehuda Tzfati, Stephane Marcand and Miki Shinohara for the gift of strains; Matt Neale for help with telomere blot quantification; Marta Grillo, Craig Gazzard, Romeela Joseph, Rebecca Leask, Alexander Wilkins, and Zher Wen Au for help with building plasmids and strains throughout the project; Pia Longhese for communication of results prior to publication. This work was supported by grants from the Medical Research Council (G0701428) and Cancer Research UK (C28567/A12720) to A.B., from the Swiss National Science Foundation (31003A_17544) and European Research Council (681-630) to P.C, and from Wellcome Investigator award No.110047/Z/15/Z to A.M.C..

## Author contributions

A.B. designed the study, wrote the manuscript, overviewed and contributed to execute all the genetic experiments, carried out some of the telomere length analysis and genotoxicity assays and analysed data. E.C. and P.C. designed and performed the biochemical experiments, and edited the manuscript. A.W.O. designed the Rad50 truncations for yeast two-hybrid assays. A.M.C contributed to design the study and edited the manuscript. C.C. did the initial sequence analysis to identify the MIN motif and edited the manuscript. M.M. performed and analysed the anti-checkpoint experiments. W.R.F. performed and analysed the NHEJ assays. C.S. performed the initial *Sc* telomere length analysis. M.A. carried out the yeast two-hybrid assays, and contributed to strain construction. F.K. performed the remaining genetic experiments with some contribution from A.B.

